# Assessment of chemotherapy-induced organ damage with ^68^Ga-labeled duramycin

**DOI:** 10.1101/630459

**Authors:** Anne Rix, Natascha Ingrid Drude, Anna Mrugalla, Ferhan Baskaya, Koon Yan Pak, Brian Gray, Hans-Jürgen Kaiser, René Hany Tolba, Eva Fiegle, Wiltrud Lederle, Felix Manuel Mottaghy, Fabian Kiessling

**Affiliations:** Institute for Experimental Molecular Imaging, Medical Faculty, RWTH Aachen University, Germany; Department of Nuclear Medicine, Medical Faculty, RWTH Aachen University, Germany; Molecular Targeting Technologies, Inc., West Chester, Pennsylvania, USA; Institute for Laboratory Animal Science, Medical Faculty, RWTH Aachen University, Germany; Department of Radiology and Nuclear Medicine, Maastricht University Medical Center (MUMC+), Maastricht, The Netherlands

**Keywords:** duramycin, apoptosis, toxicity, PET/CT, chemotherapy

## Abstract

Compared to standard toxicological techniques in preclinical toxicity studies, non-invasive imaging of organ toxicity enables fast and longitudinal investigation of the whole animal. Therefore, we set out to evaluate [^68^Ga]Ga-NODAGA-duramycin as a positron emission tomography (PET)-tracer of cell death for detecting chemotherapy-induced organ toxicity.

**Methods:** NODAGA-duramycin was radiolabeled with ^68^Ga, and quality control was done by thin layer chromatography and high performance liquid chromatography. Tracer specificity was determined in vitro by performing competitive binding experiments on ethanol treated cells. To optimize the timing of the PET/CT-based tracer evaluation, kinetic studies were performed in untreated and cisplatin-treated (20 mg/kg BW, intraperitoneal (i.p.)) BALB/cAnNRj mice. Organ uptake was analyzed in doxorubicin (4 mg/kg BW, i.p.)-, busulfan (18.8 mg/kg KG, i.p.)-, and cisplatin-treated (20 mg/kg BW, i.p.) mice, and in untreated control mice 2 hours after intravenous injection of 5–10 MBq [^68^Ga]Ga-NODAGA-duramycin. For immunofluorescence validation, tissue sections were stained with anti-active caspase-3 antibody. Blood and serum samples were collected to determine platelet count, aspartate transaminase, alanine transaminase, urea, creatinine, and creatine kinase values.

**Results:** In vitro experiments confirmed specific binding of [^68^Ga]Ga-NODAGA-duramycin to dying cells. The biodistribution analysis revealed a blood half-life of 10–17 minutes and a predominantly urinary excretion of the radiotracer. Doxorubicin-, busulfan-, and cisplatin-induced organ toxicities were detected successfully using [^68^Ga]Ga-NODAGA-duramycin PET/CT and confirmed by immunohistochemistry as well as blood parameter analysis. Busulfan-related spleno-, cardio-, and pneumotoxicity as well as cisplatin-induced cardio- and pneumotoxicity were detected even earlier by [^68^Ga]Ga-NODAGA-duramycin PET/CT than by blood parameters and histological stainings. In livers and kidneys, differences between treated and untreated animals tended to occur in PET/CT at later time points than in histology due to the relatively high background in these organs. However, trends over time were comparable.

**Conclusion:** [^68^Ga]Ga-NODAGA-duramycin PET/CT was successfully applied to non-invasively detect chemotherapy-induced organ toxicity with high sensitivity in preclinical studies. It even depicted some toxic effects prior to immunohistochemistry and blood parameter analysis and represents a promising alternative or complementary method to standard toxicological analyses. Furthermore, the tracer has a high translational potential and may provide a valuable link between preclinical and clinical research.

## Introduction

The evaluation of toxic side effects in rodents is a mandatory step in drug development prior to human trials. Current preclinical standard toxicological analyses include behavioral observations together with histopathological, biochemical, and hematological analyses. However, histopathological investigations such as hematoxylin and eosin stainings suffer from inter- and intra-observer variability, provide only limited spatial information and may not be representative of the entire organ. Furthermore, longitudinal screening is impossible, resulting in the need to sacrifice high numbers of animals [1]. Blood parameters can be obtained longitudinally and are the most applied and cost-efficient measures to evaluate health states in humans and animals. However, it is not always possible to unequivocally assign the measures to a specific organ. Furthermore, the distribution of the markers in the blood occurs with a delay after organ damage and changes may not be detected early due to their strong dilution in the blood. Finally, the amount of blood that can be collected from mice is limited, which is particularly critical in longitudinal investigations.

Today, animal experiments are strictly regulated and the 3R principles (replacement, reduction, and refinement) first described by Russell and Burch in 1959 provide an ethical framework for animal research. In addition to restricting animal experiments to the necessary minimum, the 3R principles aim to improve the quality of animals’ life and the scientific output of the experiments [2]. In this view, using non-invasive imaging techniques, like positron emission tomography (PET), for toxicity evaluation can reduce the required number of animals by enabling longitudinal investigations. Furthermore, the scientific output is improved by reducing sources of errors.

Severe organ toxicity is usually accompanied by an increased rate of cell death [3]. Depending on the underlying mechanism, cell death can be roughly divided into programmed cell death (apoptosis) and unprogrammed cell death (necrosis). During both mechanisms, the phospholipids on the inner leaflet of the cell membrane become accessible to tracers, either as a result of the decreasing cell membrane asymmetry associated with apoptosis or due to the loss of cell membrane integrity during necrosis [4]. (Pre)clinical studies focusing on in vivo cell death imaging are mostly based on molecular imaging of dying cells using labeled annexin V. Annexin V binds to phosphatidylserine, a normally inward facing phospholipid of the cell’s lipid bilayer [5]. However, annexin V tracers have some disadvantages, including a slow clearance and a limited tissue penetration, generating a considerably high unspecific background due to their high molecular weight (≥ 36 kDa) [6]. Duramycin, a peptide of 19 amino acids, is a low molecular weight alternative (2 kDa) that binds to phosphatidylethanolamine (PE), the second most abundant phospholipid on the inner leaflet of viable cells [7]. In previous studies, derivatives of duramycin have been used successfully in the detection of therapy-induced tumor cell death, in the assessment of pulmonary injury and cardiac or cerebral ischemia-reperfusion in different animal models using single-photon emission computed tomography (SPECT) [8]. Recently, Johnson et al. provided a first proof-of-concept for the suitability of ^99m^Tc-labeled duramycin to systemically characterize chemotherapy induced cell death in rats [9]. Compared to SPECT, PET provides a higher sensitivity [10]. By now, only one study has been published using duramycin as a PET tracer. Yao et al. synthesized N-(2-^18^F-fluoropropionyl)duramycin and focused on the detection of cisplatin or cyclophosphamide induced cell death, however, solely in the tumor tissue [11]. Alternatively, duramycin can be labeled using ^68^Ga [12], a generator-eluted radionuclide that combines economical and practical advantages, including no need for a cyclotron on site (as for ^18^F) and a less time-consuming synthesis due to the applicability of coordination chemistry [13]. In addition, chelating systems such as NODAGA (1-(1,3-carboxypropyl)-1,4,7-triazacyclononane-4,7-diacetic acid) allow radiolabeling at room temperature which is preferred especially for temperature sensitive peptides or proteins. Disadvantages arising from the higher maximum positron energy of ^68^Ga (1.92 MeV compared to ^18^F (0.63 MeV)), which can potentially result in a lower spatial resolution (1.4 mm (^18^F) versus 2.4 mm (^68^Ga) in a small animal PET scanner) [14] as well as the higher dose rate, need to be considered. On the other hand, the shorter half-life of ^68^Ga as compared to ^18^F can be beneficial for longitudinal studies since it allows imaging at shorter intervals.

Consequently, we evaluated [^68^Ga]Ga-NODAGA-duramycin as a ^68^Ga-based duramycin tracer for the non-invasive detection of chemotherapy-induced organ toxicity in mice using PET combined with computed tomography (CT). For this purpose, mice received three chemotherapeutic drugs with different mechanisms of action and toxicity profiles: doxorubicin, busulfan, and cisplatin. Doxorubicin is an anthracycline, which is routinely used to treat different types of cancer, including breast, lung, and thyroid cancers. Doxorubicin is especially known to cause cardiotoxic side effects, but macrophage depletion has also been reported [15, 16]. Busulfan is an alkylating agent used for the treatment of chronic myelocytic leukemia and for myeloablative conditioning prior to hematopoietic stem cell transplantations. In addition to having a myelosuppressive effect, busulfan is associated with pneumo- and cardiotoxicity [17, 18]. Cisplatin is a platinum coordination compound widely used in the treatment of various types of cancer, including cancers of soft tissue, lungs, and blood vessels. Cisplatin is known for its nephro- and neurotoxicity, but cardio-, hepato-, spleno-, and pulmotoxic effects have also been detected [19, 20]. Here we show that [^68^Ga]Ga-NODAGA-duramycin PET/CT reliably depicts organ toxicity of these chemotherapeutics in agreement with immunohistochemical analysis of caspase-3 activation and blood/serum parameters.

## Materials and methods

### Radiolabeling of NODAGA-duramycin

3.92 nmol NODAGA-duramycin (Molecular Targeting Technologies, Inc.) were radiolabeled with 40–100 MBq [^68^Ga]GaCl_3_at pH 4-5. Radiochemical yield (>95%) and purity (>95%) was estimated via thin layer chromatography and high performance liquid chromatography, respectively (for details see supplemental material and supplemental Fig. 1).

### In vitro competitive binding analysis

Binding specificity of [^68^Ga]Ga-NODAGA-duramycin was tested in vitro in triplicates using human glioblastoma *cell*s (U-87), neural stem cells (NSC), prostate cancer cells (PC-3 wt) and benign prostatic hyperplasia cells (BPH-1). To induce phosphatidylethanolamine exposure to the outer cell membrane mediated by oxidative stress, apoptosis, and necrosis, 25–30 % ethanol was added to cells with and without competitive blocking. Cells were incubated for 1h. Binding of the tracer was compared between an untreated control group, the ethanol-treated control group, and an ethanol-treated competition group. After ethanol incubation, all cells were incubated with 0.1 MBq [^68^Ga]Ga-NODAGA-duramycin for 90 minutes. The competition group was simultaneously incubated with [^68^Ga]Ga-NODAGA-duramycin and a 100-fold excess of unlabeled duramycin (40 µg/well; 90 min). After extraction of the medium, washing with PBS, and separation of dead and living cells via centrifugation, fractions of interest were measured for radioactivity using the automatic gamma counter WIZARD^2^ (Perkin Elmer). The decay corrected cell accumulated activity (dead + living) was calculated relative to the implemented activity and normalized to the overall cell number (dead and alive) as estimated with a Neubauer chamber.

### Animal model

All animal experiments were performed according to German legal requirements and animal protection laws and were approved by the Authority for Environment Conservation and Consumer Protection of the State of North Rhine-Westphalia (LANUV).

Mice were housed in groups of three to five per cage in individually ventilated cabinets under specific pathogen-free conditions with a 12 hours light- and dark-cycle in a temperature and humidity-controlled environment according to the guidelines of the “Federation for Laboratory Animal Science Associations” (www.felasa.eu). Water and standard pellets for laboratory mice (Sniff GmbH) were offered ad libitum. Female BALB/cAnNRj mice (age 10–12 weeks; Janvier Labs) were assigned randomly to four groups including untreated controls. Doxorubicin (LC Laboratories) was injected intraperitoneally (i.p.) at day 0, 7, and 14. Toxicity analyses were performed at day 8 and 15. Busulfan (Fresenius Kabi) was injected i.p. at day 0, 2, and 4. Toxicity analyses were performed at day 3 and 5. Cisplatin (Teva) was injected i.p. once. Toxicity analyses were performed 1, 2, and 3 days after injection of cisplatin. The dosages chosen for doxorubicin, busulfan, and cisplatin administration have been shown to cause toxic effects in mice in previous studies [21–23]. The toxicity analyses included [^68^Ga]Ga-NODAGA-duramycin PET/CT, immunohistochemistry, and blood/serum parameter analysis (animal numbers are listed in supplemental table 1).

### PET/CT

For PET/CT scans a 360° fly-mode CT acquisition (512 views, 3 – 5 minutes) was performed followed by a three hours-or 20 minutes-PET scan for biodistribution or toxicity analysis, respectively. Data were reconstructed using a 3-dimensional ordered-subset expectation maximization (3D-OSEM) iterative algorithm. Representative images of transversal section for visualization of lung and heart and maximum intensity projections of tracer accumulation for the figures have been created using PMOD.

PET/CT measurements were performed at 2h (except for biodistribution studies) after i.v. injection of 5–10 MBq [^68^Ga]Ga-NODAGA-duramycin using the Triumph II small animal PET/SPECT/CT (Trifoil Imaging). All data were corrected for attenuation, scatter, dead time and decay. For calculation of organ accumulated radioactivity, volumes of interest (VOI) of 0.27 mm³ were drawn using Imalytics Preclinical software [24] and their mean intensity values calculated. Additionally, a VOI was placed over the whole mouse to determine the total tracer amount. Signal intensities were converted into kBq/mL using a device-specific calibration factor. To correct for errors caused during manual injection, the amount of [^68^Ga]Ga-NODAGA-duramycin in different organs was calculated as percentage of total dose in the whole animal (TD) at the first measurement. For biodistribution analysis values were normalized to the decay corrected TD.

### Biodistribution analysis

[^68^Ga]Ga-NODAGA-duramycin biodistribution was investigated in cisplatin-treated and untreated mice (n = 3 per group). Image analysis was performed at 11, 40, 70, 100, 130, 160, and 190 minutes post-injection and time activity curves (TACs) plotted for organs of interest. The blood filled ventricles of the heart were depicted for blood concentration and blood half-lifes were calculated using a two-phase model for non-linear fitting of the TACs (GraphPad Software).

### Blood/Serum analysis

Blood and serum analysis was performed to determine platelet count (PLT), aspartate transaminase (AST), alanine transaminase (ALT), urea, creatinine, and creatine kinase (CK) values.

### Indirect immunohistochemistry

Five randomly selected vision fields per organ were analyzed (for animal numbers see supplemental table 1). Staining cryosections with 4’,6-diamidino-2-phenylindole (DAPI, Merck) and active caspase-3, which, as duramycin, is an early marker of cell damage [25], was performed as described previously using rabbit-anti-active caspase-3 IgG (6.45 μg/mL) (Abcam), followed by donkey F(ab’)2 anti-rabbit IgG (H+L)-Cy3 (3 μg/mL) (Dianova) [26]. Area percentages of active caspase-3 and DAPI per fluorescence micrograph were determined using the software Fiji [27].

### Statistical analysis

Statistical analyses were performed using GraphPad Prism 5 (GraphPad Software). Significant differences between groups were assessed using one-way ANOVA followed by Dunnett’s multiple comparisons test. *P*-values < 0.05 were considered statistically significant. All data are presented as mean ± SD.

## Results

### Competitive binding analysis

Binding specificity of [^68^Ga]Ga-NODAGA-duramycin was evaluated in vitro by competitive binding analysis using human glioblastoma cell*s* (U-87), neural stem cells (NSC), prostate cancer cell line (PC-3 wt) and benign prostatic hyperplasia cells (BPH-1). In all cell lines quantification of the decay corrected relative cell accumulated activity (rCAA), normalized to the overall cell number and relative to the implemented activity (Fig. 1) showed a significantly increased [^68^Ga]Ga-NODAGA-duramycin binding after ethanol-treatment. Competition with unlabeled duramycin significantly decreased (*p*<0.001) the uptake of [^68^Ga]Ga-NODAGA-duramycin compared to treated control cells, irrespectively of the used cell line (U-87: 24.8 ± 1.2 rCAA versus 0.5 ± 4.0 rCAA; NSC: 28.7 ± 0.4 rCAA versus 5.5 ± 2.9 rCAA; BPH-1: 26.4 ± 1.2 rCAA versus 9.3 ± 3.3 rCAA; PC-3 wt: 33.3 ± 2.9 rCAA versus 9.6 ± 1.2 rCAA).

**Figure 1.**
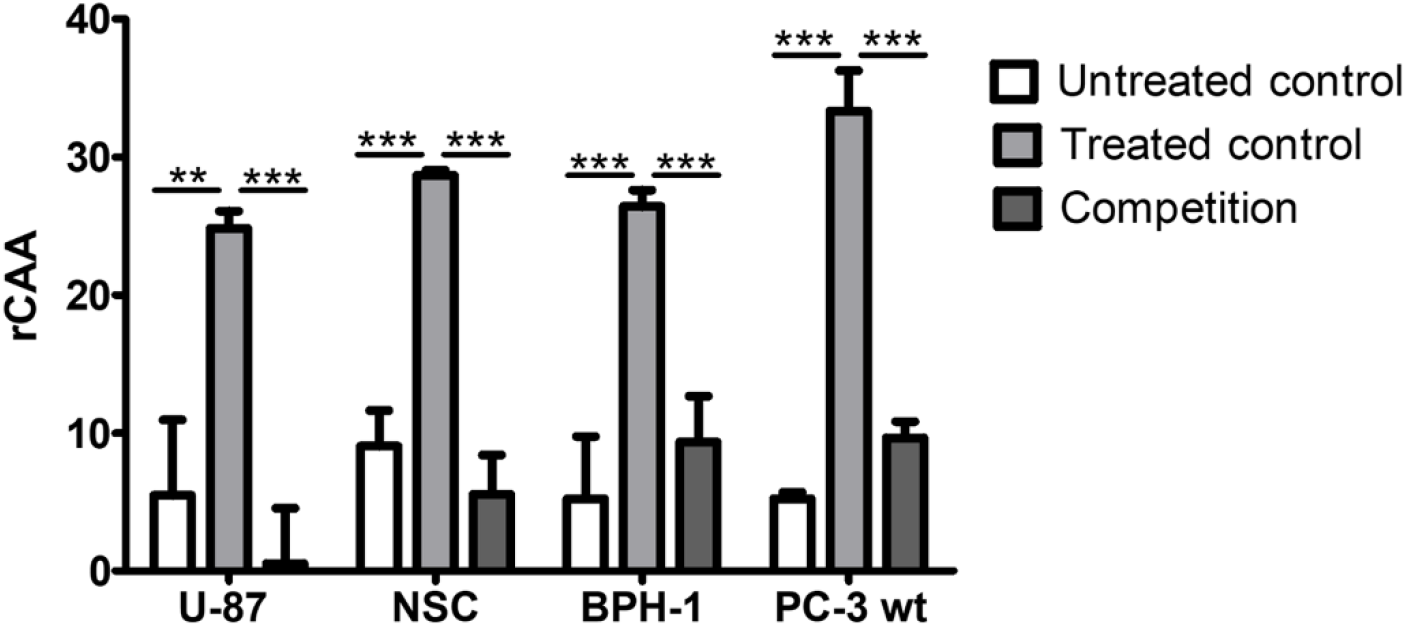
In vitro competitive binding analysis of [^68^Ga]Ga-NODAGA-duramycin in malign and benign cell lines in a cell death and stress assay using ethanol and a 100-fold excess of unlabeled duramycin (competition). Tracer uptake was significantly reduced in the competition groups. rCAA = relative cell accumulated activity; \*\**p*<0.005; \*\*\**p*<0.001.

### In vivo biodistribution

TACs of [^68^Ga]Ga-NODAGA-duramycin (Fig. 2) revealed that [^68^Ga]Ga-NODAGA-duramycin distributed rapidly throughout the body with highest initial tracer concentrations in blood and liver. Blood tracer concentration decreased rapidly, and within 120 minutes most of the unbound tracer was cleared from circulation. The blood half-life was 10.10 ± 9.25 minutes for untreated and 17.27 ± 4.12 minutes for cisplatin-treated animals. In cisplatin-treated mice, mean radioactivity in livers and spleens was consistently higher than in untreated animals and did not show a continuous decline with decreasing blood concentrations. In the kidneys an enhanced tracer accumulation became visible in cisplatin-treated animals after 70 minutes. Slightly higher values were also found in the hearts of cisplatin-treated animals, however, curves rapidly declined indicating a strong contribution of the blood to the signal at early time points. In all organs either a plateau level or a decline close to baseline was reached after 130 minutes. Radioactivity in the bladder increased until 70 – 90 minutes post injection (p.i.) indicating renal excretion.

**Figure 2.**
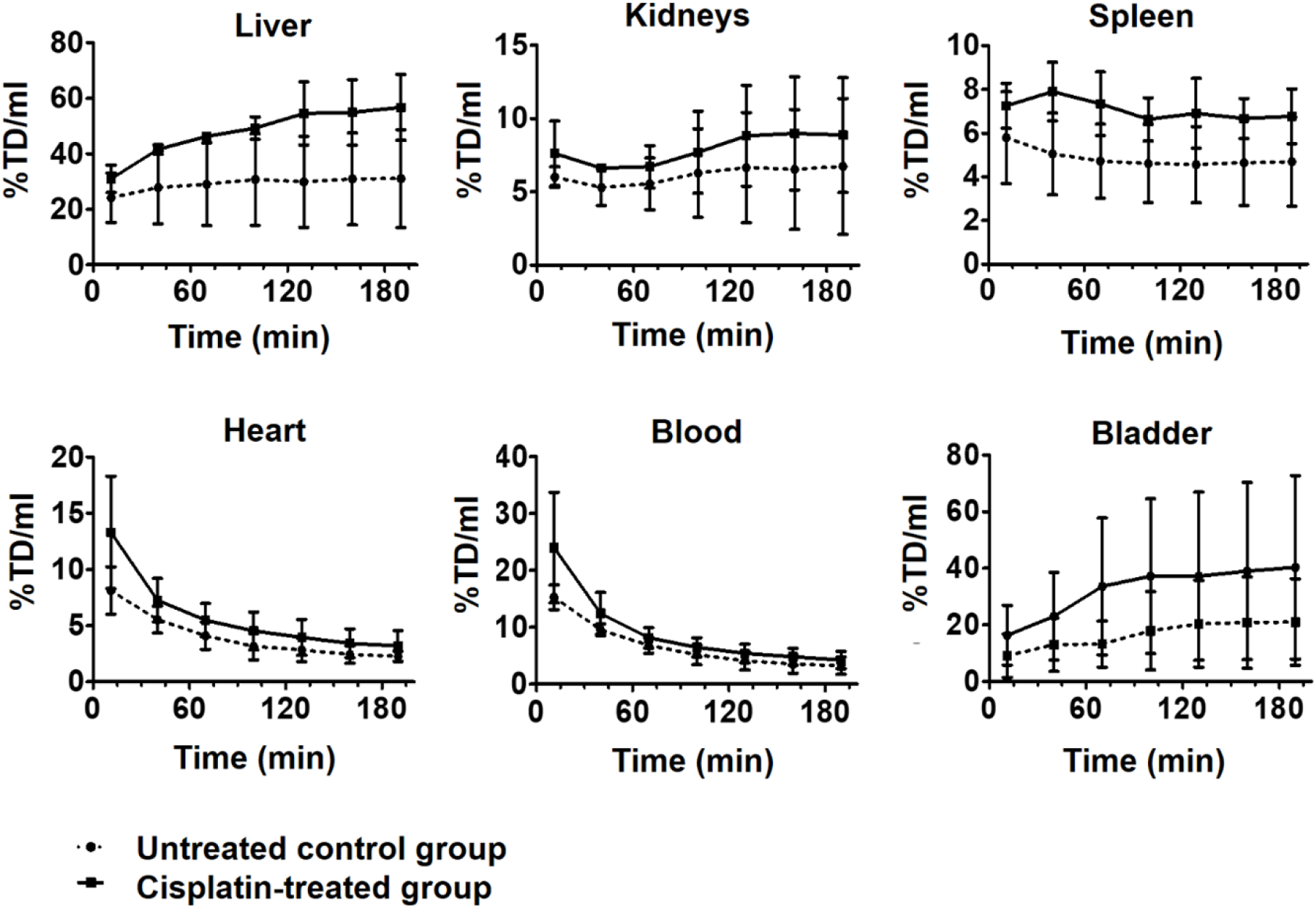
[^68^Ga]Ga-NODAGA-duramycin biodistribution displayed as TACs (10-180 min p.i. in cisplatin-treated and untreated mice. Results indicated a retention in liver and kidneys, and an accumulation in the bladder. Blood concentrations decreased rapidly. Higher tracer uptake in liver, kidneys and spleen after cisplatin treatment indicated its specificity for damaged organs. TD = total dose

### Doxorubicin-induced organ toxicity

Compared to untreated mice, doxorubicin administration lead to higher radiotracer concentrations in the liver (Fig. 3A), which was in line with caspase-3 activation measured by immunohistochemistry (Fig. 3B). In kidneys, radiotracer concentrations of doxorubicin-treated mice were higher at day 15 as well. However, for both, liver and kidney accumulation, no statistical significance was detected. In contrast, immunohistochemistry revealed significantly higher caspase-3 activation at day 15 [liver: (control: 2.0 ± 0.5 % area; d15: 3.5 ± 1.5 % area, *p* < 0.05); kidneys: (control: 0.4 ± 0.2 % area; d15: 1.1 ± 0.3 % area, *p* < 0.005)]. Additionally, liver and kidney damage were indicated by a significant increase in ALT and creatinine values at days 8 and 15, respectively (supplemental table 2 + 3). [ALT: (control: 61.78 ± 8.26 U/L; d8: 90.33 ± 19.5 U/L, *p* < 0.005); creatinine: (control: 13 ± 1.32 µmol/L; d8: 15.33 ± 1.16 µmol/L, *p* < 0.05; d15: 16.25 ± 0.96 µmol/L, *p* < 0.005).

**Figure 3.**
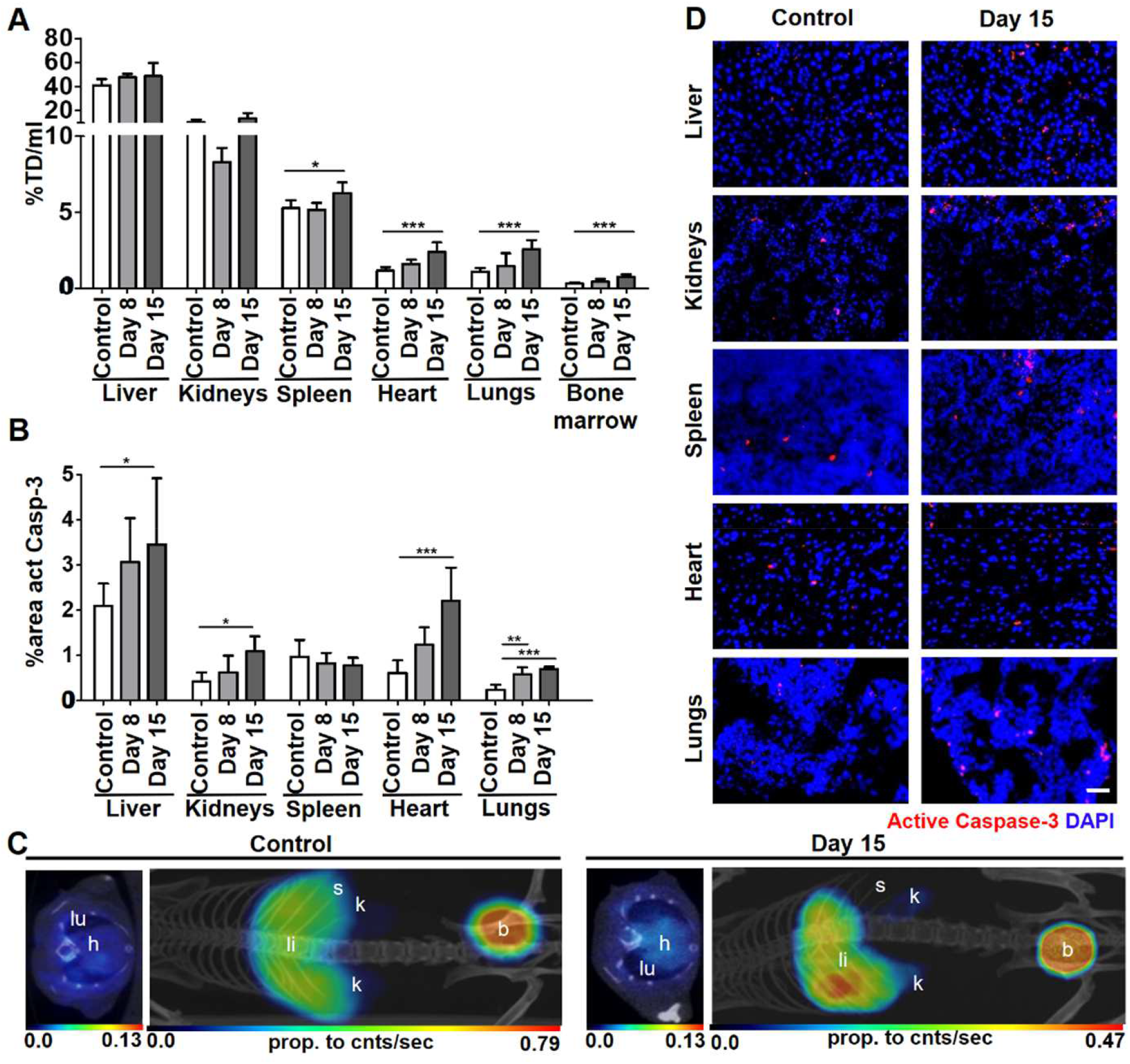
Organ toxicity analyses in doxorubicin-treated and untreated mice. (A) PET/CT showed a significantly increased accumulation in the spleen, heart, lungs and bone marrow following doxorubicin administration. (B) Caspase-3 activation revealed a significant increase in apoptosis in liver, kidneys, heart, and lungs. (C) Left: transversal section of lung and heart. Right: maximum intensity projection (MIP). (D) Representative micrographs (scale bar: 50µm). act casp-3=active caspase-3, b=bladder, d=day, h=heart, k=kidneys, li=liver, lu=lungs, s=spleen, TD=total dose; \**p*<0.05; \*\**p*<0.005; \*\*\**p*<0.001.

In spleen and bone marrow, radiotracer accumulation and blood parameters (supplemental table 3) indicated a toxic effect of doxorubicin at day 15. However, spleen toxicity was not reflected in caspase-3 activation.

In heart and lungs, all analyses indicated organ damages. Heart radiotracer accumulation in doxorubicin-treated mice was significantly increased at day 15 (control: 1.2 ± 0.2 % TD/mL; d15: 2.4 ± 0.6 % TD/mL, *p* < 0.001) accompanied by a significant increase in caspase-3 activation (control: 0.6 ± 0.3 % area; d15: 2.2 ± 0.7 % area, *p* < 0.001) and in CK levels (control: 112.4 ± 54.75 U/L; d15: 195.3 ± 34.86 U/L, *p* < 0.05). In the lungs radioactivity was increased significantly at day 15 (control: 1.1 ± 0.2 % TD/mL; d15: 2.6 ± 0.6 % TD/mL, *p* < 0.001), which corresponded to significantly higher caspase-3 activation at days 8 and 15 (control: 0.2 ± 0.1 % area; d8: 0.6 ± 0.2 % area, *p* < 0.005; d15: 0.7 ± 0.1 % area, *p* < 0.001).

### Busulfan-induced organ toxicity

Busulfan-induced hepatotoxicity was successfully detected by PET/CT (Fig. 4A) (control: 41.0 ± 5.3 % TD/mL; d5: 53.1 ± 11.8 % TD/mL; *p* < 0.005) and confirmed by immunohistochemistry (Fig. 4B) (control: 2.1 ± 0.5 % area; d5: 3.0 ± 0.1 % area, *p* < 0.005) but not by serum parameters (supplemental table 2). Nephrotoxicity analyses were consistently negative showing no significant changes in any of the applied screenings. Accumulation of [^68^Ga]Ga-NODAGA-duramycin in the bone marrow was significantly higher on days 3 and 5.

**Figure 4.**
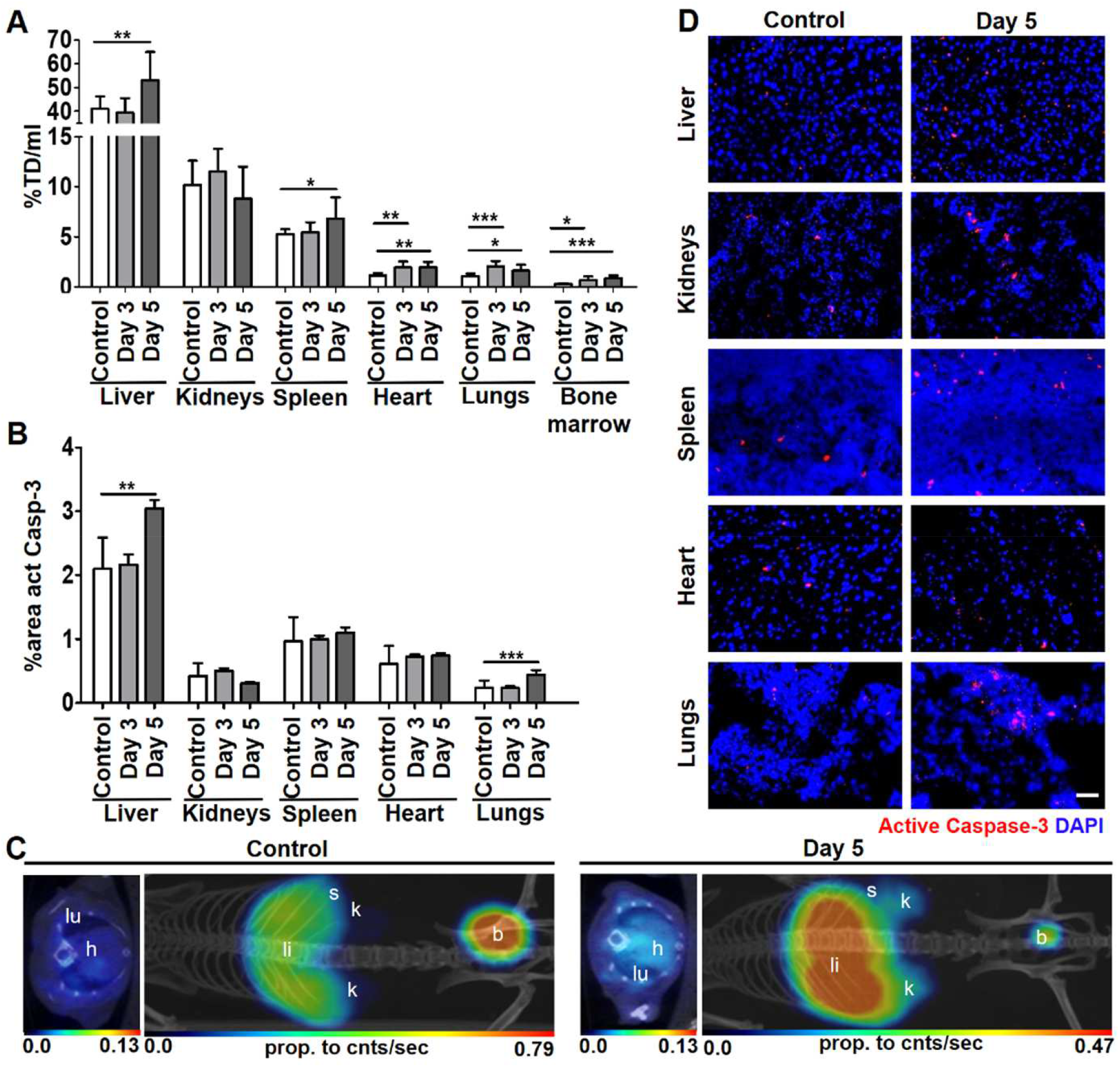
Organ toxicity analyses in busulfan-treated and untreated control mice. (A) PET/CT showed a significantly increased % TD/mL in the liver, spleen, heart, lungs, and bone marrow following busulfan administration. (B) Caspase-3 activation revealed a significant increase in apoptosis in liver and lungs. (C) Left: transversal section of lung and heart. Right: MIP. (D) Representative images (scale bar: 50µm). act casp-3=active caspase-3, b=bladder, d=day, h=heart, k=kidneys, li=liver, lu=lungs, s=spleen, TD=total dose; \**p*<0.05; \*\**p*<0.005; \*\*\**p*<0.001.

Significantly increased radioactivity in spleen and heart indicated spleno-(control: 5.3 ± 0.5 % TD/mL; d5: 6.8 ± 2.1 % TD/mL; *p* < 0.05) and cardiotoxic (control: 1.2 ± 0.2 % TD/mL; d3: 2.0 ± 0.6 % TD/mL; d5: 2.0 ± 0.5 % TD/mL; *p* < 0.005) effects. Caspase-3 activation showed a similar, however, non-significant trend with a higher variance in the control group.

In the lungs, toxicity analyses were consistent but significant differences could be detected earlier by PET/CT. In detail, significantly increased radiotracer concentrations were observed at days 3 and 5 (control: 1.1 ± 0.2 % TD/mL; d3: 2.1 ± 0.5 % TD/mL; *p <* 0.001; 1.6 ± 0.6 % TD/mL; *p* < 0.05), whereas caspase-3 activation was only increased at day 5 (control: 0.2 ± 0.1 % area; d5: 0.4 ± 0.1 % area, *p* < 0.005).

### Cisplatin-induced organ toxicity

Cisplatin-induced hepatotoxicity was detected consistently by PET/CT, immunohistochemistry and serum parameters. Compared to untreated mice, a significantly higher liver accumulation was detected at days 2 and 3 after cisplatin administration (Fig. 5A) (control: 41.0 ± 5.3 % TD/mL; d2: 54.7 ± 7.9 % TD/mL, *p* < 0.005; d3: 67.4 ± 5.8 % TD/mL, *p* < 0.001) accompanied by a significantly increased caspase-3 activation at day 3 (Fig. 5B) (control: 2.0 ± 0.5 % area; d3: 4.3 ± 0.7 % area, *p* < 0.001). Serum parameter analysis (supplemental table 2 + 3) revealed significantly higher AST levels at day 2 (control: 112.1 ± 31.75 U/L; d2: 200 ± 89.62 U/L, *p* < 0.05).

**Figure 5.**
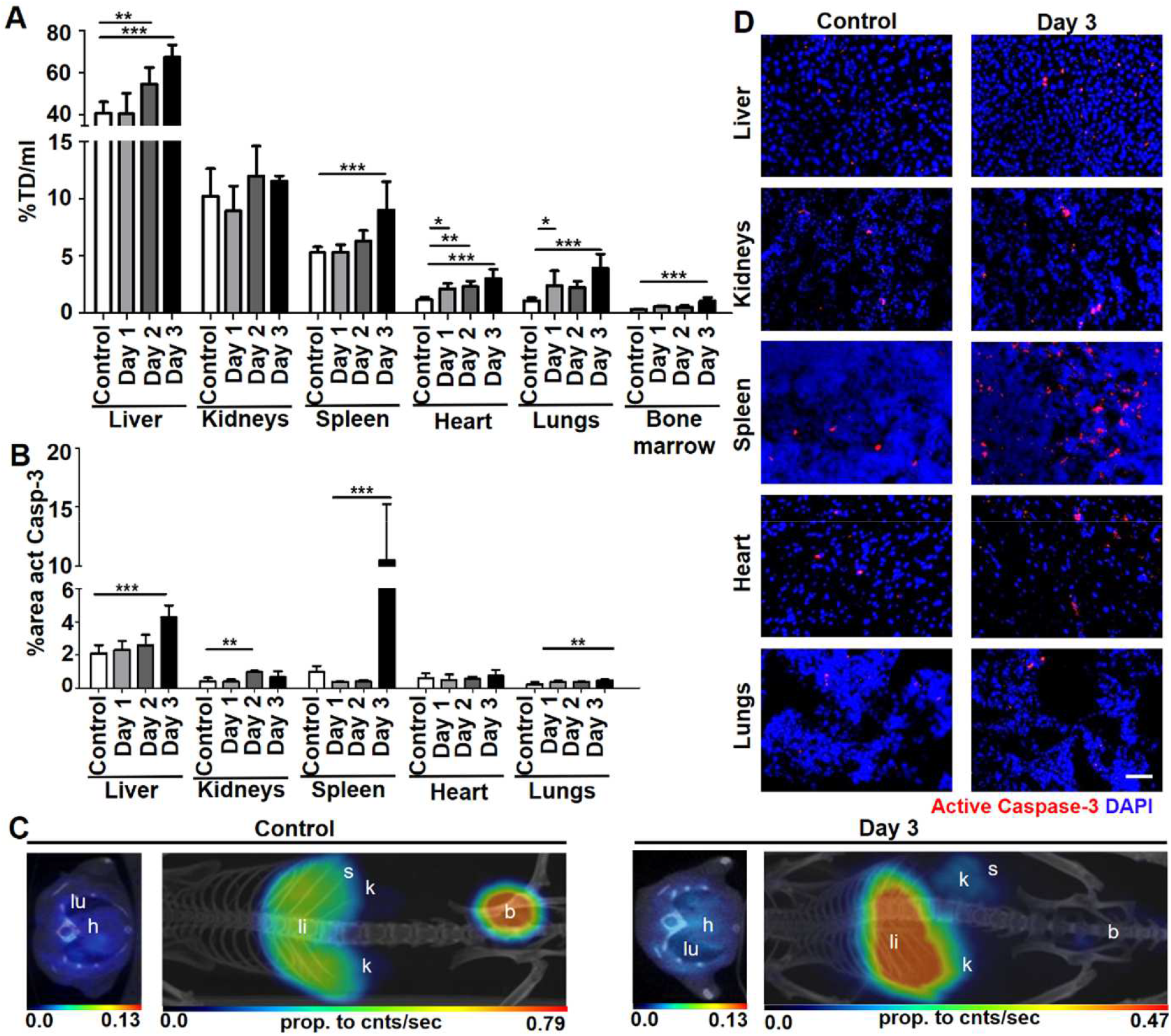
Organ toxicity analyses in cisplatin-treated and untreated control mice. (A) PET/CT images showed a significantly increased accumulation in the liver, spleen, heart, lungs, and bone marrow following cisplatin administration. (B) Caspase-3 activation showed a significant increase in apoptosis in the liver, kidneys, spleen, and lungs. (C) Left: transversal section of lung and heart. Right: MIP. (D) Representative micrographs (scale bar: 50µm). act casp-3=active caspase-3, b=bladder, d=day, h=heart, k=kidneys, li=liver, lu=lungs, s=spleen, TD=total dose; \**p*<0.05; \*\**p*<0.005; \*\*\**p*<0.001.

In the kidneys, enhanced tracer uptake was found at day 2. However, differences were not high enough to reach statistical significance. In contrast, differences in caspase-3 activation and serum parameters were more pronounced, indicating acute kidney damage at day 2 [caspase-3 activation: (control: 0.4 ± 0.2 % area; d2: 1.0 ± 0.1 % area, *p* < 0.005); urea: (control: 8.39 ± 1.23 mmol/L; d2: 15.09 ± 7.47 mmol/L, *p* < 0.05); creatinine: (control: 13 ± 1.32 µmol/L; d2: 27.67 ± 12.67 µmol/L, *p* < 0.005)].

In spleen and bone marrow, a significant increase in radiotracer accumulation was detected at day 3 compared to control animals (p < 0.001). Tracer uptake in spleen correlated with a significantly increased caspase-3 activation (control: 1.0 ± 0.2 % area; d3: 10.5 ± 4.8 % area, *p* < 0.001) and PLT (control: 894.5 ± 151.8 10³/µL; d3: 1147 ± 95.19 10³/µL, *p* < 0.005).

In the heart, radiotracer accumulation was significantly higher at all points in time (control: 1.2 ± 0.2 % TD/mL; d1: 2.1 ± 0.5 % TD/mL, *p* < 0.05; d2: 2.3 ± 0.5 % TD/mL, *p* < 0.005; d3: 3.0 ± 0.8 % TD/mL, *p <* 0.001), although no significant changes in caspase-3 activation were detected. However, significantly increased CK values at day 3 confirmed the PET/CT results (control: 112.4 ± 54.75 U/L; d3: 586.3 ± 396.8 U/L, *p* < 0.05).

In the lungs, PET/CT and immunohistochemistry were consistent, but PET/CT detected the pneumotoxic effect earlier. A significantly enhanced radiotracer accumulation indicated organ damage at days 1 and 3 after cisplatin administration (control: 1.1 ± 0.2 % TD/mL; d1: 2.4 ± 1.3 % TD/mL, d3: *p* < 0.05; 4.0 ± 1.2 % TD/mL, *p* < 0.001), while a significant increase in caspase-3 activation was only detected at day 3 (control: 0.2 ± 0.1 % area; d3: 0.5 ± 0.1 % area, *p* < 0.005).

## Discussion

The detection of drug-induced organ toxicity via non-invasive imaging could have a valuable impact on preclinical toxicity studies by improving the quality of results and reducing the number of animals needed. In this regard, cell death can be considered a suitable imaging biomarker of tissue toxicity. Recently, Johnson et al. [9] provided a proof of concept that ^99m^Tc-labeled duramycin is a suitable tracer to detect chemotherapy induced cell death by SPECT. In this study, whole body toxic profiles of cyclophosphamide, methotrexate and cisplatin were characterized on a single time point in rats. Furthermore, in a first pilot study, the progression of signal changes in rats after treatment with cyclophosphamide was detected. Based on these results, the present study aims on the longitudinal evaluation of chemotherapy induced tissue damage and the detection of systemic toxicity at an early stage using a duramycin-derivative suitable for PET imaging. Different to previous publications reporting on positron labeled duramycin [11, 12], we conjugated NODAGA to enable the complexation of ^68^Ga. Since the chelator and the radionuclide change the pharmacokinetics of duramycin [28] and since we intended to perform our analyses in mice biodistribution analyses were performed. In this context, it is noteworthy that mice generally show different pharmacokinetics and toxicity profiles than rats [29]. Consequently, we applied different chemotherapy treatment protocols to induce tissue damage in mice. Thus, although both studies confirm the suitability of duramycin as a molecule to detect cell death, the direct comparison of tracers’ performances is not possible and open to further investigation.

Specific binding of [^68^Ga]Ga-NODAGA-duramycin to damaged cells was confirmed in vitro. Analogue in vivo competitive binding experiments could not be accomplished since nephrotoxic effects of high duramycin doses in cisplatin pre-damaged kidneys inhibited renal excretion of the radiotracer and strongly altered its pharmacokinetic (data not shown). The in vivo biodistribution analysis revealed a higher blood half-life in cisplatin-treated mice compared to control animals, which might have been caused by a reduced glomerular filtration rate due to the nephrotoxicity of cisplatin. Nevertheless, the unbound radiotracer underwent fast urinary excretion. However, some retention was found in the livers and kidneys. Liver accumulation can be a result of partial biliary excretion, the higher baseline apoptosis rate in this organ, or the unspecific uptake by the mononuclear phagocyte system. Enhanced kidney concentration after 70 minutes indicated a slight unspecific kidney retention, a common observation for peptide-based tracers [30]. Both, the unspecific liver uptake and the renal retention create a baseline signal, which needs to be considered when analyzing toxicological side effects. However, since the concentration of non-specifically retained tracer was stable over time, specific accumulation could still be detected. Confirmation of small differences in tracer accumulation as valid organ toxicity may still require the complementary use of additional measures such as serum parameters [31]. Furthermore, hepatobiliary clearance and, to a lesser extent, kidney retention may be tuned by changing the chelator. However, NODAGA already appears to be one of the chelators with the lowest accumulation in the liver [32–34].

Despite the discrete unspecific uptake of [^68^Ga]Ga-NODAGA-duramycin, we were able to reliably detect doxorubicin-, busulfan- and cisplatin-induced organ toxicity by PET/CT. As expected, doxorubicin administration had a cardiotoxic effect that resulted in a significantly higher radiotracer accumulation in the heart. Additionally, toxic effects on the spleen and lungs were detected via PET/CT and confirmed by increased PLT and caspase-3 activation, respectively. This effect could be expected since doxorubicin is known to deplete CD169^+^ macrophages in the spleen and lungs [15]. Considering that splenic macrophages are responsible for the clearance of senescent platelets, an increased PLT is an additional indicator for the splenotoxic effect of doxorubicin [35].

In contrast, the hepato- and nephrotoxic effects, that were obvious in immunohistochemistry and serum parameter analysis, were less pronounced in [^68^Ga]Ga-NODAGA-duramycin PET/CT, which may be due to the enhanced unspecific uptake in these organs. As discussed above, in such cases the complementary analysis of selected serum parameters may help to confirm the PET/CT results.

The busulfan-related pneumotoxicity was detected earlier by [^68^Ga]Ga-NODAGA-duramycin PET/CT than by immunohistochemistry, which did not show a pneumotoxic effect before day 5. Additionally, organ radiotracer concentrations indicated a toxic effect on the liver, spleen, and heart. The same trend was found in immunohistochemistry, supporting previous reports in the literature [17, 18]. However, in immunohistochemistry, differences only reached significance for the hepatotoxic effect, also indicating the sensitivity of the [^68^Ga]Ga-NODAGA-duramycin PET/CT approach. The potentially higher sensitivity in the detection of toxic effects by the radiotracer can be explained by the fact that caspase-3 activation is only related to apoptosis, whereas PE becomes accessible during apoptosis, necrosis and oxidative stress [3]. Moreover, PET detects the toxicity at the whole organ level, whereas histology only focuses on a limited area of the organ. Since at early time points there are only few apoptotic cells per histological slide, the statistical validity is poor and prone to outliers. Consequently, large numbers of histological samples would need to be investigated to achieve the same sensitivity like [^68^Ga]Ga-NODAGA-duramycin PET. When comparing blood analysis with [^68^Ga]Ga-NODAGA-duramycin PET, it is worth mentioning that an elevation of blood/serum parameters often cannot be detected before substantial tissue damage occurs since molecules are diluted in the whole blood volume whereas the tracer is concentrated at a specific location in the body.

Cisplatin administration caused hepato-, spleno-, cardio- and pulmotoxic effects that were successfully detected via PET/CT and confirmed by immunohistochemistry as well as blood/serum parameter analysis, and again [^68^Ga]Ga-NODAGA-duramycin PET/CT depicted significant differences in the heart and lung prior to immunohistochemistry and serum analysis. The [^68^Ga]Ga-NODAGA-duramycin uptake in the kidneys after cisplatin treatment was in agreement with immunohistochemistry, but differences were too small and variations too high to reach statistical significance. Here we face the same situation as for doxorubicin, where an enhanced unspecific uptake makes the detection of small changes difficult. As statistical significances are depending on many factors such as number of animals per group, number of tests (multiple testing) and heterogeneity within the group, in some unclear cases it might be favorable to use each animal as its own control to detect alterations [9].

## Conclusion

[^68^Ga]Ga-NODAGA-duramycin PET/CT is a promising approach to assess drug-related side effects and organ toxicity in mice. It may be used additionally to immunohistochemistry and blood sampling and thereby, contribute to the refinement and reduction of animal experiments in pharmaceutical research. Although the analysis of serum parameters is still necessary to confirm moderate effects, [^68^Ga]Ga-NODAGA-duramycin PET/CT provides a valuable overview of all organs and reinforces the serum parameter analysis that is sometimes ambiguous regarding a specific organ. In addition, fewer serum parameters need to be analyzed, and thus, a smaller blood volume has to be collected. Furthermore, [^68^Ga]Ga-NODAGA-duramycin PET/CT allows for longitudinal investigations, which improves comparability since each animal can be used as its own control, and reduces bias in preclinical standard toxicological methods such as histopathological investigations. Beyond that, the tracer has a high translational potential that may be used to bridge the gap between preclinical and clinical research.

## Abbreviations

3R principles: replacement, reduction, and refinement
ALT: alanine transaminase
AST: aspartate transaminase
BPH-1: benign prostatic hyperplasia cells
BW: body weight
CK: creatine kinase
CT: computed tomography
DAPI: 4’,6-diamidino-2-phenylindole
i.p.: intraperitoneal
i.v.: intravenously
NODAGA: 1-(1,3-carboxypropyl)-1,4,7-triazacyclononane-4,7-diacetic acid
NSC: neural stem cells
p.i.: post injection
PC-3 wt: prostate cancer cells
PE: phosphatidylethanolamine
PET: positron emission tomography
PLT: platelet count
rCAA: relative cell accumulated activity
SD: standard deviation
SPECT: single-photon emission computed tomography
TAC: time activity curve
TD: total dose
U-87: human glioblastoma cells
VOI: volume of interest

## Acknowledgements

This study was supported by BMBF (German Federal Ministry of Education and Research) project LiSyM (Grant No. 031L0041) and by DFG (German Research Foundation) research group FOR2591 (Grant No. 321137804).

## Contributions

Anne Rix, Natascha Drude and Anna Mrugalla, wrote the manuscript. Anne Rix and Anna Mrugalla performed the experiments and acquired most of the data. Natascha Drude performed the radiolabeling and participated in the PET/CT experiments and data analysis. Ferhan Baskaya performed and analyzed the immunohistochemistry. Koon Y. Pak and Brian Gray developed and synthesized NODAGA-duramycin. Hans-Jürgen Kaiser and René Tolba provided support in animal handling, PET-CT imaging and data analysis. Eva Fiegle, Felix Mottaghy and Fabian Kiessling participated in study design and data analysis. Wiltrud Lederle, Felix Mottaghy and Fabian Kiessling provided supervisory support and edited the manuscript.

