## Supplementary data for "Assessment of chemotherapy-induced organ damage with ^68^Ga-labeled duramycin"

**MATERIALS AND METHODS**

**Radiolabeling of NODAGA-duramycin**

For radiolabeling of NODAGA-duramycin (Molecular Targeting Technologies, Inc.), no-carrier-added [^68^Ga]GaCl_3_ (in 0.05 M HCl) was eluted from an in clinic [^68^Ge]/[^68^Ga] generator (ITM Isotopen Technologien München). NODAGA-duramycin was dissolved in ultrapure water (Merck) and 3 M NH_4_CH_3_O_2(aq)_ (in ultrapure H_2_O) was added to adjust the pH to 3 – 4 for radiolabeling with ^68^Ga at room temperature. The radiochemical yield (rcy) was determined by Radio-TLC with instant Thin Layer Chromatography (iTLC)-silica strips and citrate buffer as mobile phase and was reproducibly >95%. Radiochemical purity was estimated via high performance liquid chromatography (HPLC) on a C-18 column (phenomenex) with a mobile phase linear gradient from 90 – 10 % 0.1 % TFA (in HPLC-H_2_O) with 10 – 90 % acetonitrile over 15 min, with a flow rate of 1 mL/min at room temperature (rcp > 95%).

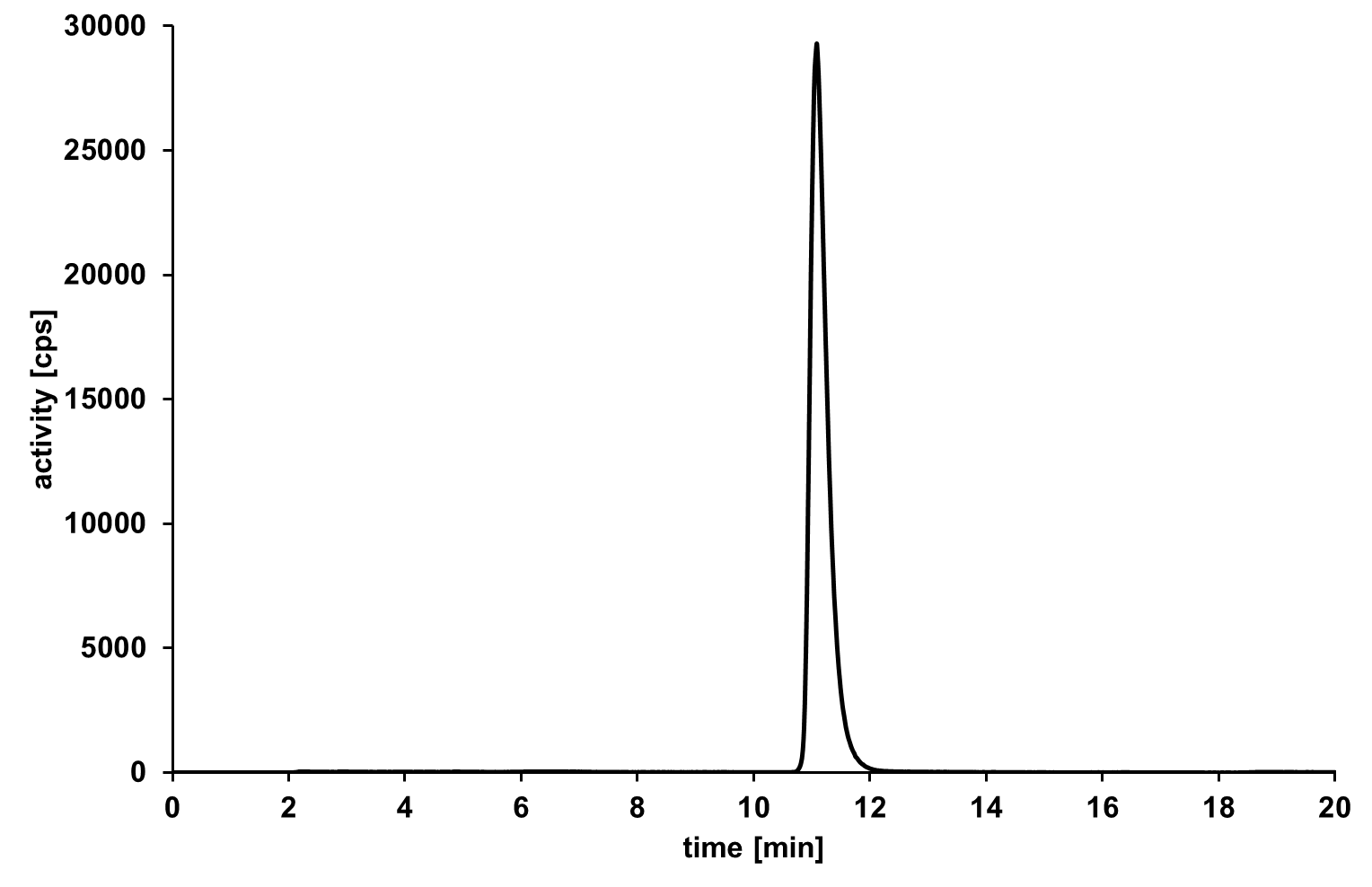

**SUPPLEMENTAL Figure 1:** Representative HPLC of ^68^Ga-labeled NODAGA-duramycin showing a radiochemical purity >95%.

**SUPPLEMENTAL TABLE 1.** List of animal numbers per group. Numbers in square brackets indicate exceptions of parameters where data had to be excluded. ALT=alanine transaminase, AST=aspartate transaminase, CK=creatine kinase.

|  | **PET/CT** | **Immunohistology** | **Blood/serum parameters** |
| --- | --- | --- | --- |
| **control** | 9 | 9 [8 (heart), 6 (spleen)] | 13 [10 (urea, creatinine, CK, ALT, and AST)] |
| **doxorubicin d8** | 4 | 3 | 3 |
| **doxorubicin d15** | 5 | 4 [3 (liver and lungs)] | 4 |
| **busulfan d3** | 10 | 3 | 6 |
| **busulfan d5** | 9 | 5 | 11 [10 (urea, creatinine, CK, ALT, and AST)] |
| **cisplatin d1** | 4 | 3 | 3 |
| **cisplatin d2** | 5 | 3 | 12 [9 (urea, creatinine, CK), 8 (ALT, AST)] |
| **cisplatin d3** | 5 | 4 [3 (liver and lungs)] | 5 [4 (urea, creatinine, CK, ALT, and AST)] |

**SUPPLEMENTAL TABLE 2.** Serum parameter analysis of control, doxorubicin-, busulfan- and cisplatin-treated mice showed a significant increase and PLT levels after cisplatin administration. **p*<0.05; ***p*<0.005; ****p*<0.001.

|  | **AST** | **ALT** | **Urea** | **Creatinine** | **CK** |
| --- | --- | --- | --- | --- | --- |
|  | **(U/L)** | **(U/L)** | **(mmol/L)** | **(µmol/L)** | **(U/L)** |
| **Control** | 112.1 | 61.78 | 8.39 | 13 | 112.4 |
|  | (+/−31.75) | (+/−8.26) | (+/−1.23) | (+/−1.32) | (+/−54.75) |
| **Doxorubicin** | 94.67 | 90.33** | 7.73 | 15.33* | 109.7 |
| **d8** | (+/−9.29) | (+/−19.5) | (+/−0.175) | (+/−1.16) | (+/−29.14) |
|  | 96.5 | 52.25 | 7 | 16.25** | 195.3* |
| **d15** | (+/−9.98) | (+/−7.41) | (+/−0.46) | (+/−0.96) | (+/−34.86) |
| **Busulfan** | 82.5 | 55.33 | 7.81 | 11.17 | 141.5 |
| **d3** | (+/−29.66) | (+/−14.73) | (+/−1.25) | (+/−1.17) | (+/−60.39) |
|  | 84.4 | 87.1 | 7.19 | 12.6 | 160.2 |
| **d5** | (+/−33.46) | (+/−84.03) | (+/−1.22) | (+/−2.5) | (+/−81.33) |
| **Cisplatin** | 122 | 58 | 6.32 | 16.67 | 180.7 |
| **d1** | (+/−18.33) | (+/−2) | (+/−0.24) | (+/−1.53) | (+/−44.19) |
|  | 200* | 88.88 | 15.09* | 27.67** | 372.6 |
| **d2** | (+/−89.62) | (+/−49.2) | (+/−7.47) | (+/−12.67) | (+/−255.2) |
|  | 183.8 | 63.5 | 11.08 | 22.25 | 586.3* |
| **d3** | (+/−59.73) | (+/−20.79) | (+/−1.37) | (+/−5.44) | (+/−396.8) |

**SUPPLEMENTAL TABLE 3.** Blood parameter analysis of control, doxorubicin-, busulfan- and cisplatin-treated mice showed a significant decrease in the red blood cell count (RBC), hemoglobin (HGB) and hematocrit (HCT), and a significant increase in the number of platelets (PLT) after doxorubicin administration. Furthermore, busulfan and cisplatin significantly decreased the white blood cell counts (WBC). **p*<0.05; ***p*<0.005; ****p*<0.001.

|  | **RBC** | **HGB** | **HCT** | **WBC** | | **PLT** |
| --- | --- | --- | --- | --- | --- | --- |
|  | **(10^6^/µl)** | **(g/dL)** | **(%)** | **(10^3^/µl)** | | **(10^3^/µl)** |
| **Control** | 9.22 | 14.6 | 39.8 | 6.83 | | 894.5 |
|  | (+/−0.36) | (+/−0.6) | (+/−1.5) | (+/−1.4) | | (+/−151.8) |
| **Doxorubicin** | 9.62 | 14.8 | 41.2 | 9.2 | | 1011 |
| **d8** | (+/−0.09) | (+/−0.1) | (+/−0.2) | (+/−2.0) | | (+/−22.5) |
|  | 7.98*** | 12.6*** | 34.1*** | 6.5 | | 1106* |
| **d15** | (+/−0.58) | (+/−1.2) | (+/−2.8) | (+/−2.3) | | (+/−51.55) |
| **Busulfan** | 9.79 | 15.2 | 42.0 | 4.73* | | 849.3 |
| **d3** | (+/−0.38) | (+/−0.4) | (+/−1.7) | (+/−1.48) | | (+/−82.9) |
|  | 8.32 | 13.3 | 35.2 | 4.94* | | 1175* |
| **d5** | (+/−2.02) | (+/−3.3) | (+/−8.5) | (+/−1.61) | | (+/−531.1) |
| **Cisplatin** | 10.23 | 15.9* | 43.5** | 5.63 | | 797 |
| **d1** | (+/−0.25) | (+/−0.1) | (+/−0.3) | (+/−0.55) | | (+/−19.67) |
|  | 9.45 | 15.08 | 40.5 | 4.85** | 919 | |
| **d2** | (+/−0.49) | (+/−0.85) | (+/−2.3) | (+/−1.69) | (+/−134.5) | |
|  | 9.81* | 15.38 | 41.6 | 3.68** | 1147** | |
| **d3** | (+/−0.29) | (+/−0.6) | (+/−1.6) | (+/−0.97) | (+/−95.19) | |
